# Multimodal brain age estimates relate to Alzheimer disease biomarkers and cognition in early stages: a cross-sectional observational study

**DOI:** 10.1101/2022.08.25.505251

**Authors:** Peter R Millar, Brian A Gordon, Patrick H Luckett, Tammie LS Benzinger, Carlos Cruchaga, Anne M Fagan, Jason J Hassenstab, Richard J Perrin, Suzanne E Schindler, Ricardo F Allegri, Gregory S Day, Martin R Farlow, Hiroshi Mori, Georg Nübling, the Dominantly Inherited Alzheimer Network, Randall J Bateman, John C Morris, Beau M Ances

## Abstract

**Background:** Estimates of “brain-predicted age” quantify apparent brain age compared to normative trajectories of neuroimaging features. The brain age gap (BAG) between predicted and chronological age is elevated in symptomatic Alzheimer disease (AD), but has not been well explored in preclinical AD. Prior studies have typically modeled BAG with structural magnetic resonance imaging (MRI), but more recently other modalities, including functional connectivity (FC) and multimodal MRI, have been explored.

**Methods:** We trained three models to predict age from FC, volumetric (Vol), or multimodal MRI (Vol+FC) in 390 control participants (18-89 years old). In independent samples of 144 older adult controls, 154 preclinical AD participants, and 154 cognitively impaired (CI; CDR > 0) participants, we tested relationships between BAG and AD biomarkers of amyloid, tau, and neurodegeneration, as well as a global cognitive composite.

**Results:** All models predicted age in the control training set, with the multimodal model outperforming the unimodal models. All three BAG estimates were significantly elevated in CI compared to controls. FC-BAG and Vol+FC-BAG were marginally reduced in preclinical AD participants compared to controls. In CI participants only, elevated Vol-BAG and Vol+FC-BAG were associated with more advanced AD pathology and lower cognitive performance.

**Conclusions:** Both FC-BAG and Vol-BAG are elevated in CI participants. However, FC and volumetric MRI also capture complementary signals. Specifically, FC-BAG may capture a unique biphasic response to preclinical AD pathology, while Vol-BAG may capture pathological progression and cognitive decline in the symptomatic stage. A multimodal age-prediction model captures these modality-specific patterns, and further, improves sensitivity to healthy age differences.

**Funding:** This work was supported by the National Institutes of Health (P01-AG026276, P01-AG03991, P30-AG066444, 5-R01-AG052550, 5-R01-AG057680, 1-R01-AG067505, 1S10RR022984-01A1, U19-AG032438), the BrightFocus Foundation (A2022014F), and the Alzheimer’s Association (SG-20-690363-DIAN).

## Introduction

Alzheimer disease (AD) is marked by structural and functional disruptions in the brain, some of which can be observed through multimodal magnetic resonance imaging (MRI) in preclinical and symptomatic stages of the disease^1,2^. More recently, the “brain-predicted age” framework has emerged as a promising tool for neuroimaging analyses, leveraging recent developments and accessibility of machine learning techniques, as well as large-scale, publicly available neuroimaging datasets^3,4^. These models are trained to quantify how “old” a brain appears, as compared to a normative sample of training data - typically consisting of cognitively normal participants across the adult lifespan (e.g., 5). Thus, the framework allows for a residual-based interpretation of the brain age gap (BAG), defined as the difference between model-predicted age and chronological age, as an index of vulnerability and/or resistance to underlying disease pathology. Indeed, several studies have demonstrated that BAG is elevated (i.e., the brain “appears older” than expected) in a host of neurological and psychiatric disorders, including symptomatic AD^6,7,8^, as well as schizophrenia (e.g., 9), HIV (e.g., 10), and type-2 diabetes (e.g., 11), and moreover, predicts mortality (12). Conversely, lower BAG is associated with lower risk of disease progression^6,13,14^. Critically, at least one comparison suggests that BAG exceeds other established MRI (hippocampal volume) and CSF (pTau, Aβ42) biomarkers in sensitivity to AD progression (6). Thus, by summarizing complex, non-linear, highly multivariate patterns of neuroimaging features into a simple, interpretable summary metric, BAG may reflect a comprehensive biomarker of brain health.

Several studies have established that symptomatic AD and mild cognitive impairment (MCI) are associated with elevated BAG^3,4^. However, the sensitivity of these model estimates to AD in the preclinical stage (i.e., present amyloid pathology in the absence of cognitive decline^15^) is less clear. The development of sensitive, reliable, non-invasive biomarkers of preclinical AD pathology is critical for the assessment of individual AD risk, as well as the evaluation of AD clinical prevention trials. Recent studies have demonstrated that greater BAG is associated with greater amyloid PET burden in a Down syndrome cohort (16), and with greater tau PET burden in sporadic MCI and symptomatic AD (17). Another model attempted to improve sensitivity to preclinical sporadic AD, by training only on participants without evidence of amyloid pathology (18). Compared to a typical amyloid-insensitive model (5), the restricted model appeared to be more sensitive to progressive stages of AD (18). However, this comparison included amyloid-negative and amyloid-positive test samples from two separate cohorts, and thus may be driven by cohort, scanner, and/or site differences. To validate the applicability of the brain-predicted age approach to preclinical AD, it is important to test a model’s sensitivity to amyloid status, as well as continuous relationships with preclinical AD biomarkers, within a single cohort.

Most of the brain-predicted age reports described above focused primarily on structural MRI. However, other studies have successfully modeled brain age using a variety of other modalities, including metabolic PET (17,19), diffusion MRI (20,21), and functional connectivity (FC)^22–25^. Integration of multiple neuroimaging modalities may maximize sensitivity of BAG estimates to preclinical AD. Indeed, recent multimodal comparisons suggest that structural MRI and FC capture complementary age-related signals (24,26) and that age prediction may be improved by incorporating multiple modalities (25,27). One recent study has shown that BAG estimates from a FC graph theory-based model are significantly elevated in autosomal dominant AD mutation carriers and are positively associated with amyloid PET (28). Further, we have recently demonstrated that FC correlation-based BAG estimates are surprisingly reduced in preclinical sporadic AD participants with evidence of amyloid pathology and elevated pTau, as well as in cognitively normal APOE ε4 carriers at genetic risk of AD (29). Thus, incorporating FC into BAG models may improve sensitivity to early AD.

This project aimed to develop and validate multimodal models of brain-predicted age, incorporating both FC and structural MRI. The influence of preclinical AD pathology was strictly controlled in the training set to maximize sensitivity. We hypothesized that BAG estimates would be sensitive to preclinical AD and early cognitive impairment. We further considered whether estimates were continuously associated with AD biomarkers of amyloid, tau, and neurodegeneration (30), as well as cognitive function. Finally, we systematically compared the performance of models trained on unimodal FC, structural MRI, and combined modalities, to test the added utility of multimodal integration.

## Methods

### Participants

We formed a training sample of healthy controls spanning the adult lifespan by combining structural and FC-MRI data from three sources, as described previously (29): the Charles F. and Joanne Knight Alzheimer Disease Research Center (ADRC) at Washington University in St.

Louis (WUSTL), healthy controls from studies in the Ances lab at WUSTL (31,32), and mutation-negative controls from the Dominantly Inherited Alzheimer Network (DIAN) study of autosomal dominant AD at multiple international sites including WUSTL (33). To minimize the likelihood of undetected AD pathology in our training set, participants over the age of 50 were only included in the training set if they were cognitively normal (CN), as assessed by the Clinical Dementia Rating^®^ (CDR^®^ 0) (34), and had at least one biomarker indicating the absence of amyloid pathology (A-, see below). We excluded 59 participants who did not have available CDR or biomarker measures (see Figure S1). As CDR and amyloid biomarkers were not available in the Ances lab controls, we included only participants at or below age 50 from this cohort in the training set. These healthy control participants were randomly divided into a training set (~80%; N = 390) and a held-out test set (~20%; N = 97), which did not significantly differ in age, sex, education, or race, see Table 1.

**Table 1.**
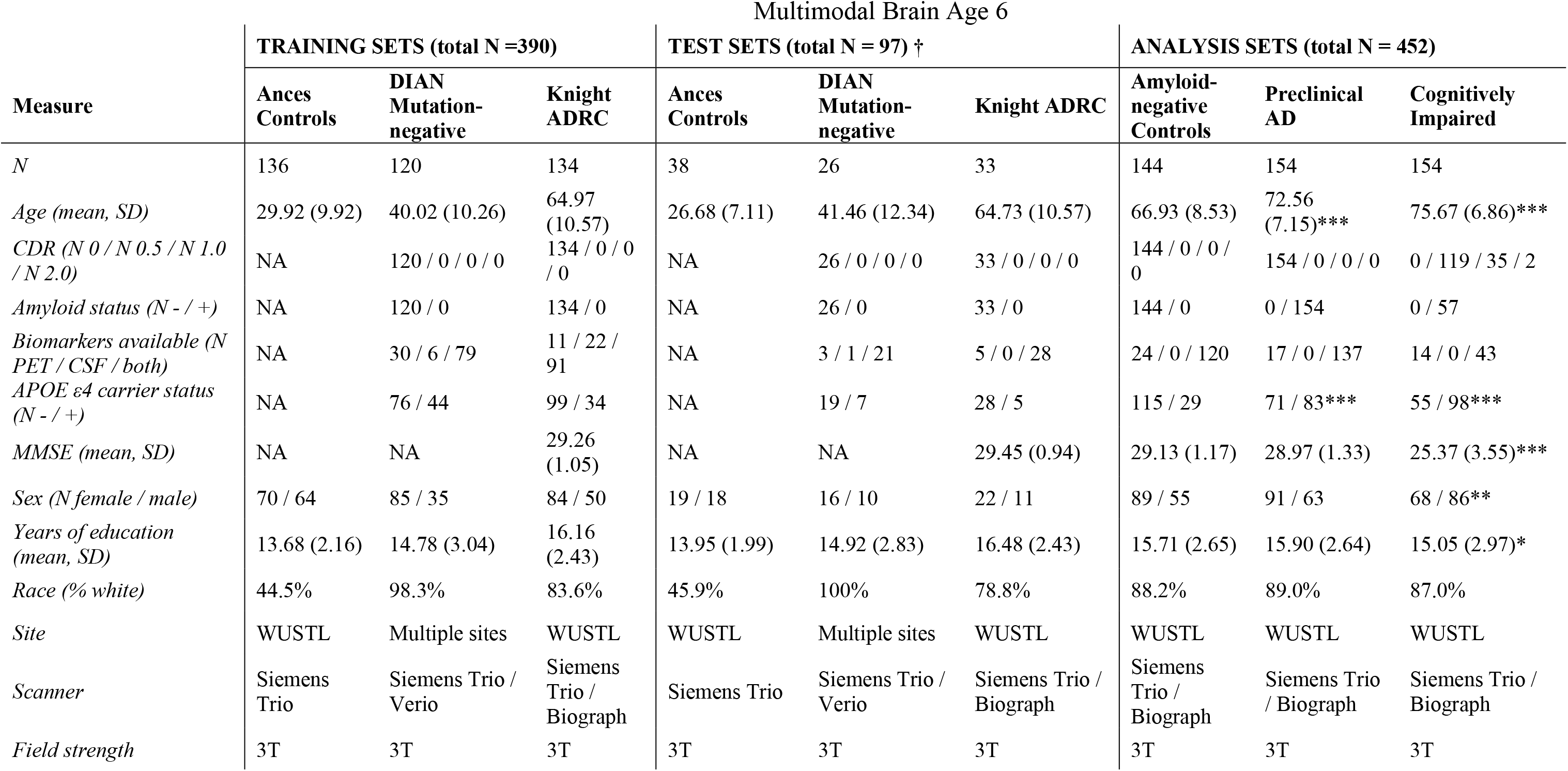
Demographic information of the combined samples. DIAN = Dominantly Inherited Alzheimer Network, ADRC = Alzheimer Disease Research Center, AD = Alzheimer disease, CDR = Clinical Dementia Rating, MMSE = Mini Mental State Examination, WUSTL = Washington University in St. Louis, T = Tesla. Group differences from the amyloid-negative controls were tested with *t* tests for continuous variables and *χ2* tests for categorical variables. \*\*\**p* < .001, ** *p* < .01, * *p* < .05, ^*p* < .10. † Note: Test sets include randomly-selected, non-overlapping subsets of participants drawn from the same studies as the training sets.

Finally, independent samples for hypothesis testing included three groups from the Knight ADRC: a randomly selected sample of 144 CN, A-controls who did not overlap with the training or testing sets, 154 preclinical AD participants (CN, A+), and 154 cognitively impaired (CI) participants (CDR > 0 with a biomarker measure consistent with amyloid pathology (see below) and/or a primary diagnosis of AD or uncertain dementia (35)). See Table 1 for demographic details of each sample. All procedures were approved by the Human Research Protection Office at WUSTL.

### PET & CSF Biomarkers

Amyloid burden was imaged with positron emission tomography (PET) using [11C]-Pittsburgh Compound B (PIB) (36) or [18F]-Florbetapir (AV45) (37). Regional standard uptake ratios (SUVRs) were modeled from 30 to 60 minutes after injection for PIB and from 50 to 70 minutes for AV45, using cerebellar grey as the reference region (38). Regions of interest were segmented automatically using FreeSurfer 5.3 (39). Global amyloid burden was defined as the mean of partial-volume-corrected (PVC) SUVRs from bilateral precuneus, superior and rostral middle frontal, lateral and medial orbitofrontal, and superior and middle temporal regions (38). Amyloid summary SUVRs were harmonized across tracers using a Centiloid conversion (40).

Tau deposition was imaged with PET using [18F]-Flortaucipir (AV-1451) (41). Regional SUVRs were modeled from 80 to 100 minutes after injection, using cerebellar grey as the reference region. A tau summary measure was defined in the mean PVC SUVRs from bilateral amygdala, entorhinal, inferior temporal, and lateral occipital regions (42).

Cerebrospinal fluid (CSF) was collected via lumbar puncture using methods described previously (43). After overnight fasting, 20- to 30-mL samples of CSF were collected, centrifuged, then aliquoted (500 μL) in polypropylene tubes, and stored at −80°C. CSF amyloid β peptide 42 (Aβ42), Aβ40, and phosphorylated tau-181 (pTau) were measured with automated Lumipulse immunoassays (Fujirebio, Malvern, PA) using a single lot of assays for each analyte. Aβ42 and pTau estimates were each normalized for individual differences in CSF production rates by forming a ratio with Aβ40 as the denominator (44,45). CSF neurofilament light (NfL) was measured with an ELISA immunoassay (Uman Diagnostics, Umeå, Sweden). As both pTau/Aβ40 and NfL were highly skewed, we applied a log transformation to these estimates before statistical analysis.

Amyloid positivity was defined using previously published cutoffs for PIB (SUVR > 1.42) (46) or AV45 (SUVR > 1.19) (47). Additionally, the CSF Aβ42/Aβ40 ratio has been shown to be highly concordant with amyloid PET (positivity cutoff < 0.0673) (48,49). Thus, participants were defined as amyloid-positive (for preclinical AD and CI groups) if they had either a PIB, AV45, or CSF Aβ42/Aβ40 ratio measure in the positive range. Participants with discordant positivity between PET and CSF estimates were defined as amyloid-positive. We used a Gaussian mixture model approach to define pTau positivity based on the CSF pTau/Aβ40 ratio.

### Cognitive Battery

Knight ADRC participants completed a 2-hour battery of cognitive tests. We examined global cognition by forming a composite of tasks across cognitive domains, including processing speed (Trail Making A)^50^, executive function (Trail Making B)^50^, semantic fluency (Animal Naming)^51^, and episodic memory (Free and Cued Selective Reminding Test free recall score)^52^. This composite has recently been used to study individual differences in cognition in relation the preclinical AD biomarkers and structural MRI (53), as well as functional MRI measures (54).

### MRI Acquisition

All MRI data were obtained using a Siemens 3T scanner, although there was a variety of specific models within and across studies. As described previously (29), participants in the Knight ADRC and Ances lab studies completed one of two comparable structural MRI protocols, varying by scanner (sagittal T1-weighted magnetization-prepared rapid gradient echo sequence [MPRAGE] with repetition time [TR] = 2400 or 2300 ms, echo time [TE] = 3.16 or 2.95 ms, flip angle = 8° or 9°, frames = 176, field of view = sagittal 256×256 or 240×256 mm, 1-mm isotropic or 1×1×1.2 mm voxels; oblique T2-weighted fast spin echo sequence [FSE] with TR = 3200 ms, TE = 455 ms, 256 x 256 acquisition matrix, 1-mm isotropic voxels) and an identical resting-state fMRI protocol (interleaved whole-brain echo planar imaging sequence [EPI] with TR = 2200 ms, TE = 27 ms, flip angle = 90°, field of view = 256 mm, 4-mm isotropic voxels for two 6-minute runs [164 volumes each] of eyes open fixation). There was variability in the MPRAGE and EPI sequence parameters for the DIAN participants (33) with the most notable difference being shorter resting-state runs (one 5-minute run of 120 volumes).

### FC Preprocessing and Features

All MRI data were processed using common pipelines. Initial fMRI preprocessing followed conventional methods, as described previously (29,55), including frame alignment, debanding, rigid body transformation, bias field correction, and normalization of within-run intensity values to a whole-brain mode of 1000 (56). Transformation to an in-house atlas template based on 120 independent, CN older adults was performed using a composition of affine transforms connecting the functional volumes with the T2-weighted and MPRAGE images. Frame alignment was included in a single resampling that generated a volumetric timeseries of the concatenated runs in isotropic 3-mm atlas space.

As described previously (29,57), additional processing was performed to allow for nuisance variable regression. Data underwent framewise censoring based on motion estimates (framewise displacement [FD] > 0.3mm and/or derivative of variance [DVARS] > 2.5 above participant’s mean). To further minimize the confounding influence of head motion on FC estimates (56) in all samples, we only included scans with low head motion (mean FD < 0.30 mm and > 50% frames retained after motion censoring). BOLD data underwent a temporal band-pass filter (0.005 Hz <*f* < 0.1 Hz) and nuisance variable regression, including motion parameters, timeseries from FreeSurfer 5.3-defined (39) whole brain (global signal), CSF, ventricle, and white matter masks, as well as the derivatives of these signals. Finally, BOLD data were spatially blurred (6 mm FWHM).

Final BOLD timeseries data were averaged across voxels within a set of 300 spherical regions of interest (ROIs) in cortical, subcortical, and cerebellar areas (58). For each scan, we calculated the 300 x 300 Fisher-transformed Pearson correlation matrix of the final averaged BOLD timeseries between all ROIs. We then used the vectorized upper triangle of each correlation matrix (excluding auto-correlations; 44,850 total correlations) as input features for predicting age. Since site and/or scanner differences between samples might confound neuroimaging estimates, we harmonized FC matrices using an empirical Bayes modeling approach (ComBat) (59,60), which has previously been applied to FC data (61).

### Structural MRI Processing and Features

All T1-weighted images underwent cortical reconstruction and volumetric segmentation through a common pipeline with FreeSurfer 5.3 (39,62). Initial processing included motion correction, segmentation of subcortical white matter and deep grey matter, intensity normalization, registration to a spherical atlas, and parcellation of the cerebral cortex based on the Desikan atlas (63). Inclusion and exclusion errors of parcellation and segmentation were identified and edited by a centralized team of trained research technicians according to standardized criteria (38). We then used the FreeSurfer-defined thickness estimates from 68 cortical regions (63), along with volume estimates from 33 subcortical regions (62) as input features for predicting age. We harmonized volumetric features across sites and scanners using the same ComBat approach (59,60), which has also been applied to volumetric data (64).

### Gaussian Process Regression (GPR)

As described previously (29), machine learning analyses were conducted using the Regression Learner application in Matlab 2021a (65). We trained two Gaussian process regression (GPR)^66^ models, each with a rational quadratic kernel function to predict chronological age using fully-processed, harmonized MRI features (FC or volumetric) in the training set. The *σ* hyperparameter was tuned within each model by searching a range of values from 10^-4^ to 10\**SD_age_* using Bayesian optimization across 100 training evaluations. The optimal value of *σ* for each model was found (see Figure S2), and was applied for all subsequent applications of that model. All other hyperparameters were set to default values (basis function = constant, standardize = true).

Model performance in the training set was assessed using 10-fold cross validation via the mean absolute error (*MAE*) between true chronological age and the cross-validated age predictions merged across the 10 folds. We then evaluated generalizability of the models to predict age in unseen data by applying the trained models to the held-out test set of healthy controls. To construct an empirical distribution of performance estimates in each model, we repeated this process in 1000 bootstrap simulations, randomly reassigning the training and held-out test data in each simulation. Finally, we applied the same fully-trained GPR models to separate analysis sets of 154 CI, 154 preclinical AD, and 144 CN, A-controls to test our hypotheses regarding AD-related group effects and individual difference relationships. Unimodal models were each constructed with a single GPR model. The multimodal model was constructed by taking the “stacked” predictions from each first-level unimodal model as features for training a second-level GPR model^25,26,27^.

For each participant, we calculated model-specific BAG estimates as the difference between chronological age and age predictions from the unimodal FC model (FC-BAG), volumetric model (Vol-BAG), and multimodal model (Vol+FC-BAG). To correct for regression dilution commonly observed in similar models (67–69), we included chronological age as a covariate in all statistical tests of BAG (16,67). However, to avoid inflating estimates of prediction accuracy (70), only *uncorrected* age prediction values were used for evaluating model performance in the training and test sets.

### Statistical Analysis

All statistical analyses were conducted in R 4.0.2 (R Core Team, 2020). Demographic differences in the AD samples were tested with independent-samples *t* tests for continuous variables and *χ^2^* tests for categorical variables, using CN, A-controls as a reference group. We tested AD group differences in BAG estimates, as well as continuous relationships with AD biomarkers and cognitive estimates using linear regression models. Since the range of amyloid biomarkers was drastically reduced in the CN A-sample, we excluded these participants from models testing continuous amyloid relationships. To correct for age-related bias in BAG^67^ (as previously mentioned), as well as demographic differences between samples, we controlled for age, sex, years of education, and race as covariates during statistical tests. Effect sizes were computed as partial *η^2^* (*η_p_^2^*). Group differences in each BAG estimate were tested using an omnibus Kruskal-Wallis test with follow-up pairwise Wilcoxon rank sum tests on age-residualized BAG estimates.

### Data availability

This project utilized datasets obtained from the Knight ADRC and DIAN. The Knight ADRC and DIAN encourage and facilitate research by current and new investigators, and thus, the data and code are available to all qualified researchers after appropriate review. Requests for access to the data used in this study may be placed to the Knight ADRC Leadership Committee (https://knightadrc.wustl.edu/professionals-clinicians/request-center-resources/) and the DIAN Steering Committee (https://dian.wustl.edu/our-research/for-investigators/dian-observational-study-investigator-resources/data-request-form/). Code used in this study is available at https://github.com/peterrmillar/MultimodalBrainAge.

## Results

### Sample Description & Demographics

Demographic characteristics of the training sets, test sets, and analysis sets are reported in Table 1. Preclinical AD participants were older (*t* = 6.15, *p* < .001) and more likely to be *APOE* ε4 carriers (*χ^2^* = 34.73, *p* < .001) than amyloid-negative controls. Further, CI participants were older (*t* = 9.71, *p* < .001), more likely male (*χ^2^* = 8.60, *p* = .003), more likely to be *APOE* ε4 carriers (*χ^2^* = 56.67,*p* < .001), and had fewer years of education (*t* = 2.03,*p* < .043), and lower MMSE scores (*t* = 12.46, *p* < .001) than amyloid-negative controls.

### Comparison of Model Performance

All models accurately predicted chronological age in the training sets, as assessed using 10-fold cross validation (*MAE_FC_* = 8.672 years, *MAE_Vol_* = 5.973 years, *MAE_Vol+FC_* = 5.340 years, see Figure 1A-C). Across 1000 random reassignments of training and test samples, model performance in held-out test data was comparable to performance in the training set (see Figure 1D). Further, in the bootstrapped held-out test samples, *MAEs* from the volumetric model were significantly lower than the FC model (*t* = 95.60,*p* < .001), while the multimodal model significantly outperformed both unimodal models (*t*’s > 37, *p*’s < .001). There was a significant, but modestly sized, positive correlation between FC-BAG and Vol-BAG in the adult lifespan CN training and testing sets (*r* = .095, *p* = .036, see Figure S3A), as well as the AD analysis sets (*r* = .134, *p* = .004, see Figure S3B).

**Figure 1.**
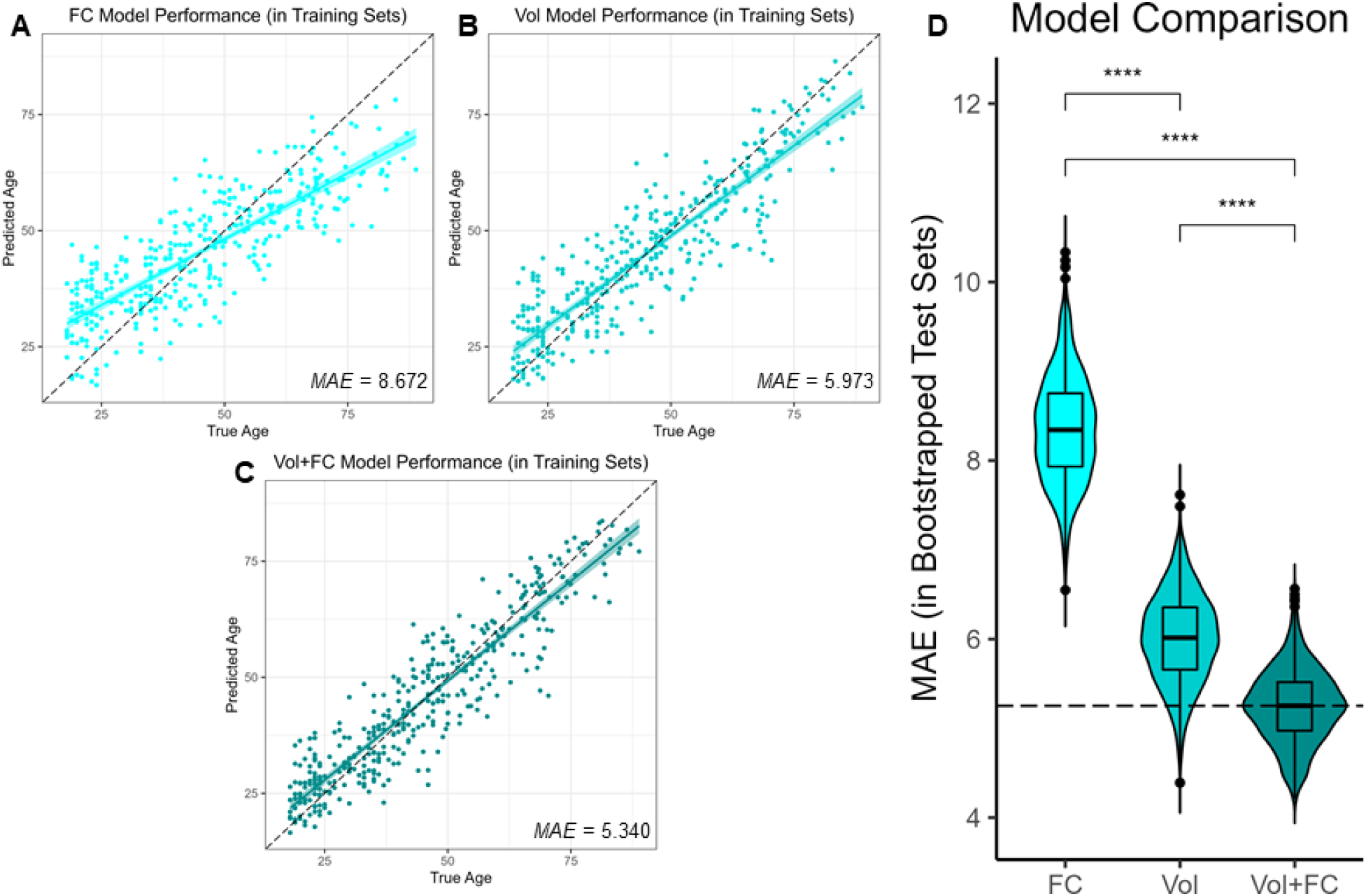
Performance of the brain age models. Scatterplots show the cross-validated model predictions in the training sets for functional connectivity (FC; A), volumetric (Vol; B) and multimodal models (Vol+FC; C). Age predicted by each model (y axis) is plotted against true age (x axis). Colored lines and shaded areas represent regression lines and 95% confidence regions. Dashed black lines represent perfect prediction. Model performance is evaluated by mean absolute error (*MAE*). Violin plot (D) shows the bootstrapped age prediction performance (*MAE*) for each model in held-out test sets across 1000 random reassignments of training and test data. Dashed black line represents median test-set performance in the best-performing (Vol+FC) model.

### BAG Differences in Cognitive Impairment and Preclinical AD

Three linear regression models tested the effects of cognitive impairment (CDR > 0 vs. CDR 0) and AD pathology (amyloid positivity [A- vs. A+] and pTau positivity [T- vs. T+]) on FC-BAG, Vol-BAG, and Vol+FC-BAG, controlling for true age and demographic covariates (see Table 2). An omnibus Kruskal-Wallis test revealed significant differences in residual FC-BAG across the four groups (*H* = 19.94, *p* < .001). FC-BAG was 3.56 years older in CI participants compared to CN controls (*β* = 3.56, *p* = 0.002, *η_p_^2^* = 0.03, see Figure 2A&B, Table 2A). Follow-up Wilcoxon tests revealed that residual FC-BAG was significantly elevated in CI relative to CN A+T- (*p* = 0.006) and A+T+ participants (*p* < 0.001). FC-BAG was also marginally lower by 1.75 years in T+ participants compared to T- (*β* = −1.75, *p* = 0.088, *η_p_^2^* = 0.01), but did not significantly differ as a function of amyloid positivity, controlling for global CDR and the other predictors. Follow-up Wilcoxon tests revealed that residual FC-BAG showed a stepwise decline with increasing preclinical AD pathology, such that compared to CN A-T-controls, FC-BAG was significantly lower in CN A+T- (*p* = .042), as well as CN A+T+ participants (*p* < 0.001), and marginally lower in A+T+ participants compared to A+T- (*p* = 0.110).

**Figure 2.**
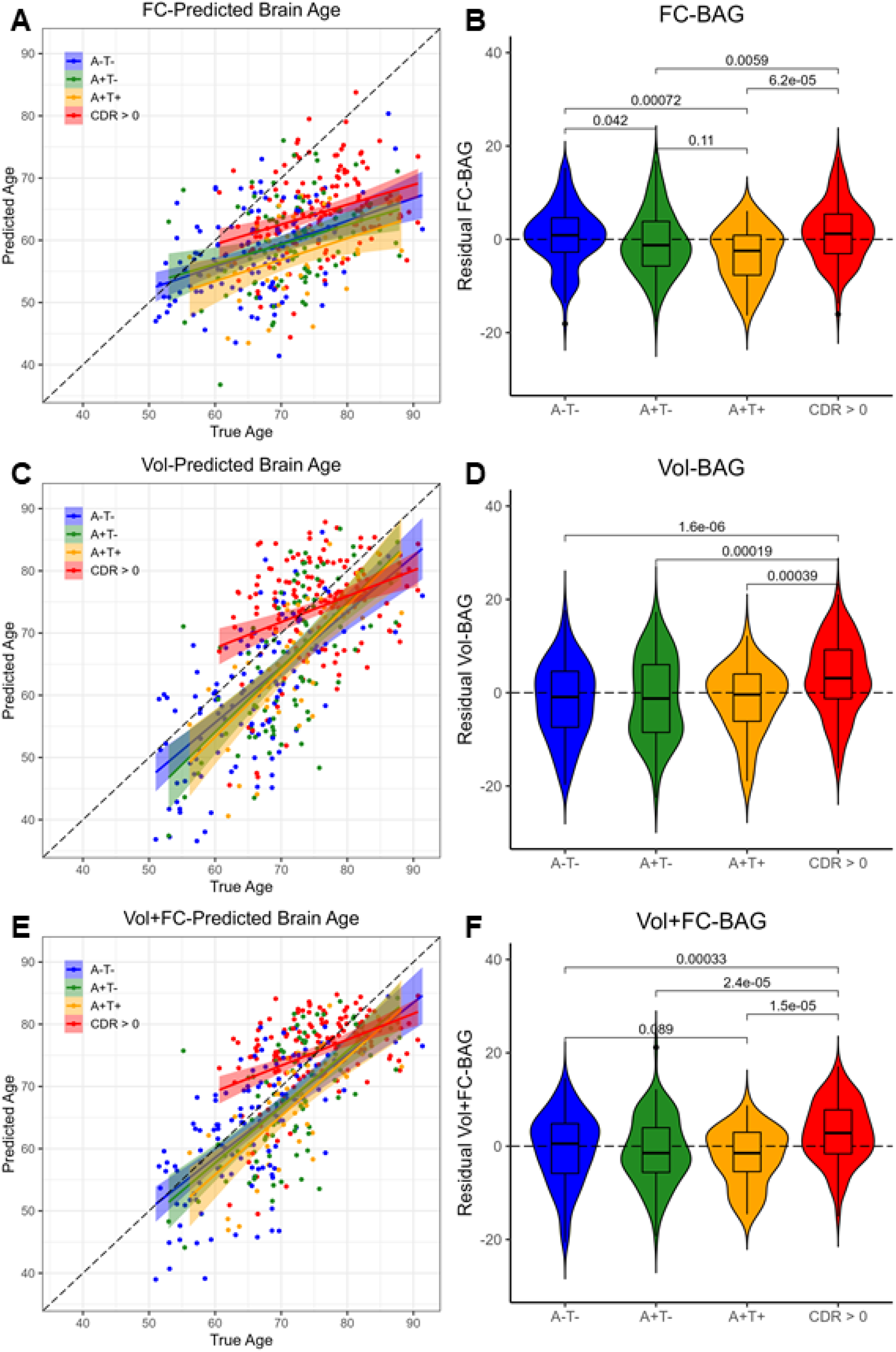
Group differences in functional connectivity (FC; A, B), volumetric (Vol; C, D), and multimodal (Vol+FC; E, F) brain age in the analysis sets. Comparisons are presented between cognitively normal (CDR = 0) biomarker-negative controls (A-T-; blue) vs. A+T- (green) vs. A+T+ (gold) vs. cognitively impaired participants (CDR > 0, red). Scatterplots (A, C, E) show predicted vs. true age for each group. Colored lines and shaded areas represent group-specific regression lines and 95% confidence regions. Dashed black lines represent perfect prediction. Violin plots (B, D, F) show residual FC-BAG (controlling for true age) in each group. P values are reported from pairwise independent-samples Wilcoxon rank sum tests.

**Table 2.** Linear regression models predicting FC-BAG (A), Vol-BAG (B), and FC+Vol-BAG (C). Model estimates are presented as beta weight (standard error). CDR = Clinical Dementia Rating. *** p < .001, ** p < .01, * p < .05, ^ p < .10.

| *Dependent Variable* | A. FC-BAG | B. Vol-BAG | C. Vol+FC-BAG |
| --- | --- | --- | --- |
| **Coefficients** |  |  |  |
| *Intercept* | 35.14 (4.13)*** | 2.15 (5.53) | 12.16 (4.75)* |
| *CDR > 0* | 3.56 (1.14)** | 4.73 (1.52)** | 4.61 (1.31)*** |
| *Amyloid +* | -0.29 (0.89) | 0.68 (1.22) | -0.18 (1.02) |
| *Tau +* | -1.75 (1.02)^ | -1.55 (1.19) | -0.26 (1.18) |
| *Age (y)* | -0.65 (0.05)*** | -0.1 (0.07) | -0.19 (0.05)*** |
| *Sex = female* | -1.29 (0.72)^ | 1.44 (0.98) | 0.65 (0.83) |
| *Education (y)* | 0.01 (0.13) | -0.23 (0.18) | -0.25 (0.15)^ |
| *Race = white* | 2.5 (1.34)^ | 2.37 (1.97) | 2.26 (1.54) |

A second omnibus Kruskal-Wallis test revealed significant differences in residual Vol-BAG across the four groups (*H* = 25.81, *p* < .001). Vol-BAG was 4.73 years older in CI participants compared to CN controls (*β* = 4.73, *p* = 0.002, *η_p_^2^* = 0.03, see Figure 2C&D, Table 2B). Follow-up Wilcoxon tests revealed that residual Vol-BAG was significantly elevated in CI participants relative to CN A-T-, A+T-, and A+T+ participants (p’s < 0.001). Vol-BAG did not significantly differ as a function of amyloid or tau positivity, controlling for CDR and the other covariates.

A third omnibus Kruskal-Wallis test revealed significant differences in residual Vol+FC-BAG across the four groups (H = 27.21, p < .001). Vol+FC-BAG was 4.61 years older in CI participants compared to CN controls (β = 4.61, p < 0.001, η_p_^2^ = 0.04, see Figure 2E&F, Table 2C). Follow-up Wilcoxon tests revealed that residual FC-BAG was significantly elevated in CI participants relative to CN A-T-, A+T-, and A+T+ participants (p’s < 0.001). Vol+FC-BAG did not significantly differ as a function of amyloid or tau positivity, controlling for CDR and the other predictors. However, follow-up Wilcoxon tests revealed that residual Vol+FC-BAG was marginally lower in CN A+T+ relative to A-T-participants (p = 0.089).

### Relationships with Amyloid Markers

In the functional connectivity model, FC-BAG was not significantly related with amyloid PET, nor was there an interactive relationship with amyloid PET between groups (see Figure 3A). There were also no significant main effects or interactions between FC-BAG, Vol-BAG, or Vol+FC-BAG and CSF Aβ42/40 (See Figure 3B, 3D, 3F).

**Figure 3.**
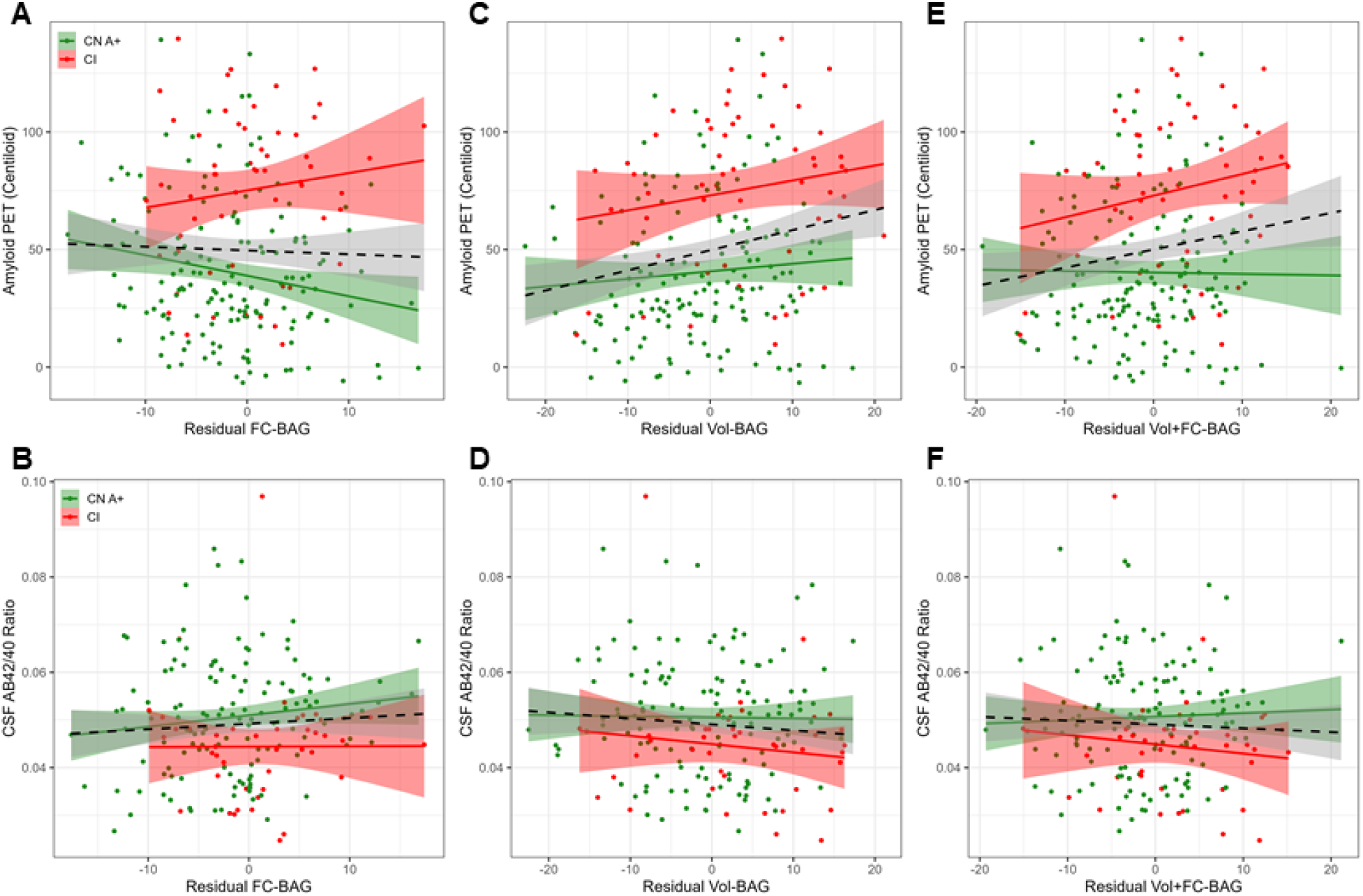
Continuous relationships between amyloid biomarkers and functional connectivity (FC-BAG; A, B), volumetric (Vol-BAG; C, D), and multimodal (Vol+FC-BAG; E, F) brain age gap in the analysis sets. Scatterplots show Amyloid PET (A, C, E) and CSF AB42/40 (B, D, F) as a function of residual BAG (controlling for true age) in each group. Colored lines and shaded areas represent group-specific regression lines and 95% confidence regions. Dashed black lines represent main effect regression lines across all groups.

In the volumetric and multimodal models, there were significant main effects, such that greater Vol-BAG (*β* = 0.79, *p* = .004, *η_p_^2^* = 0.041, see Figure 3C) and greater Vol+FC-BAG (*β* = 0.85, *p* = .011, *η_p_^2^* = 0.031, see Figure 3E) were both associated with greater amyloid PET. In the multimodal model only, this relationship was further characterized by a marginal interaction, such that the association was more strongly positive in CI participants than in preclinical AD (*β* = 1.18, *p* = .081, *η_p_^2^* = 0.015).

### Relationships with Tau Markers

In the functional connectivity model, FC-BAG was not significantly related with tau PET or CSF pTau-181/Aβ40 (see Figure 4A, 4B). However, there was a marginal interaction, suggesting a trend toward higher CSF pTau-181/Aβ40 in CI participants with high FC-BAG, but not in CN controls (*β* = 0.02, *p* = .067, *η_p_^2^* = 0.015).

**Figure 4.**
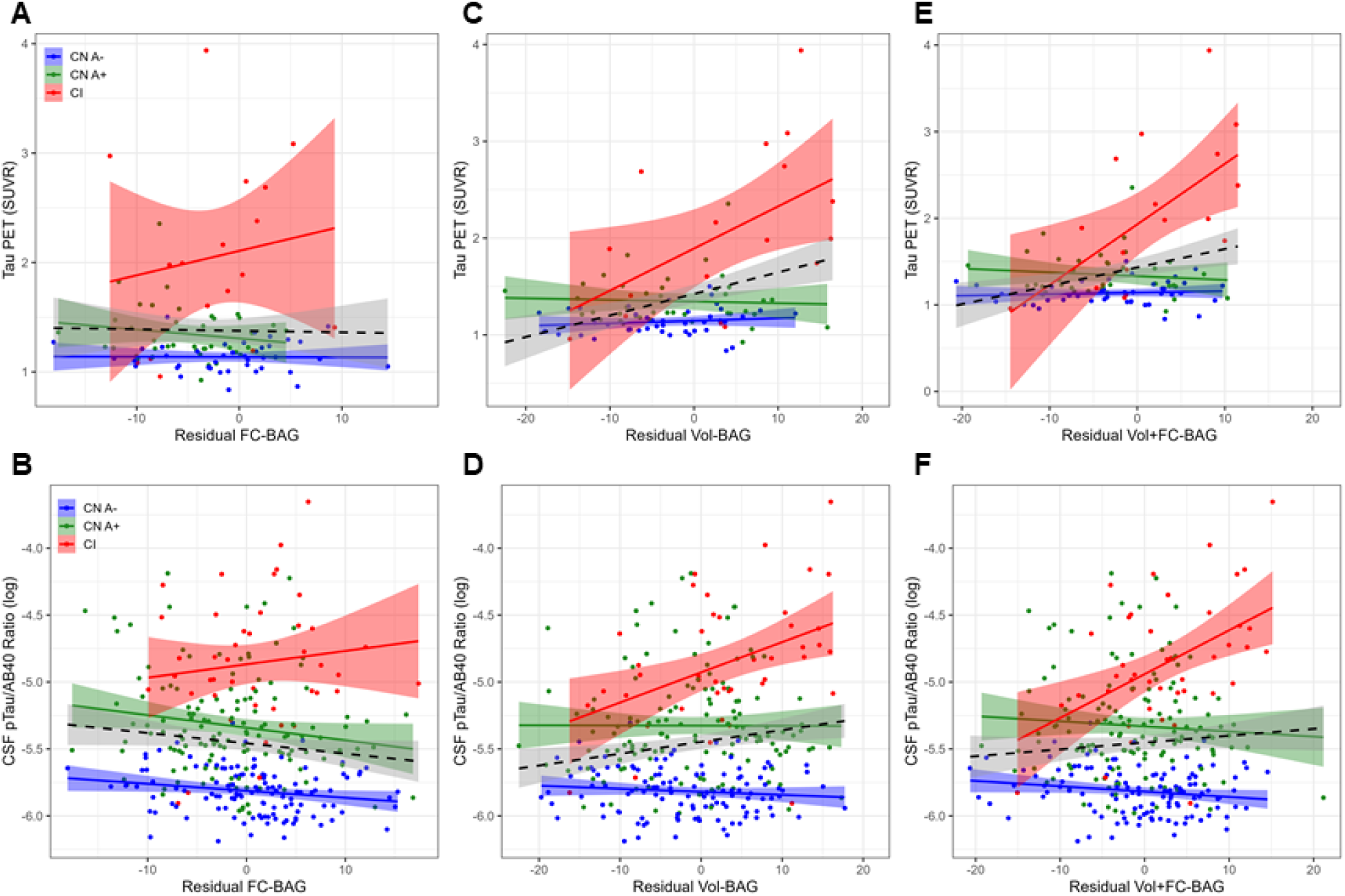
Continuous relationships between tau biomarkers and functional connectivity (FC-BAG; A, B), volumetric (Vol-BAG; C, D), and multimodal (Vol+FC-BAG; E, F) brain age gap in the analysis sets. Scatterplots show Tau PET summary (A, C, E) and log-transformed CSF pTau/Aβ40 (B, D, F) as a function of residual BAG (controlling for true age) in each group. Colored lines and shaded areas represent group-specific regression lines and 95% confidence regions. Dashed black lines represent main effect regression lines across all groups.

In the volumetric and multimodal models, there were significant main effects, such that greater Vol-BAG (*β* = 0.02, *p* < .001, *η_p_^2^* = 0.141, see Figure 4C) and greater Vol+FC-BAG (*β* = 0.02, *p* = .001, *η_p_^2^* = 0.118, see Figure 4E) were both associated with greater tau PET. These main effects were further characterized by significant interactions, such that the positive association was only observed in CI participants, but not in the other groups (Vol-BAG: *β* = 0.04, *p* < .001, *η_p_^2^* = 0.176; Vol+FC-BAG: *β* = 0.07, *p* < .001, *η_p_^2^* = 0.254).

Consistent with tau PET, CSF pTau/Aβ40 demonstrated similar interactive effects, such that greater Vol-BAG (*β* = 0.02, *p* < .001, *η_p_^2^* = 0.052, see Figure 4D) and greater Vol+FC-BAG (*β* = 0.04,*p* < .001, *η_p_^2^* = 0.073, see Figure 4F) were both associated with greater CSF pTau/Aβ40 in the CI participants, but not in the other groups.

### Relationships with Neurodegeneration Markers

In the functional connectivity model, there was a significant main effect relationship, such that across all groups, greater FC-BAG was associated with lower hippocampal volume (*β* = −23.6, *p* = .002, *η_p_^2^* = 0.022, see Figure 5A), however this effect was likely driven by group differences in both measures, as there were no significant relationships within or interactions between the three groups. There was no significant main effect or interaction relationship between FC-BAG and CSF NfL (see Figure 5B).

**Figure 5.**
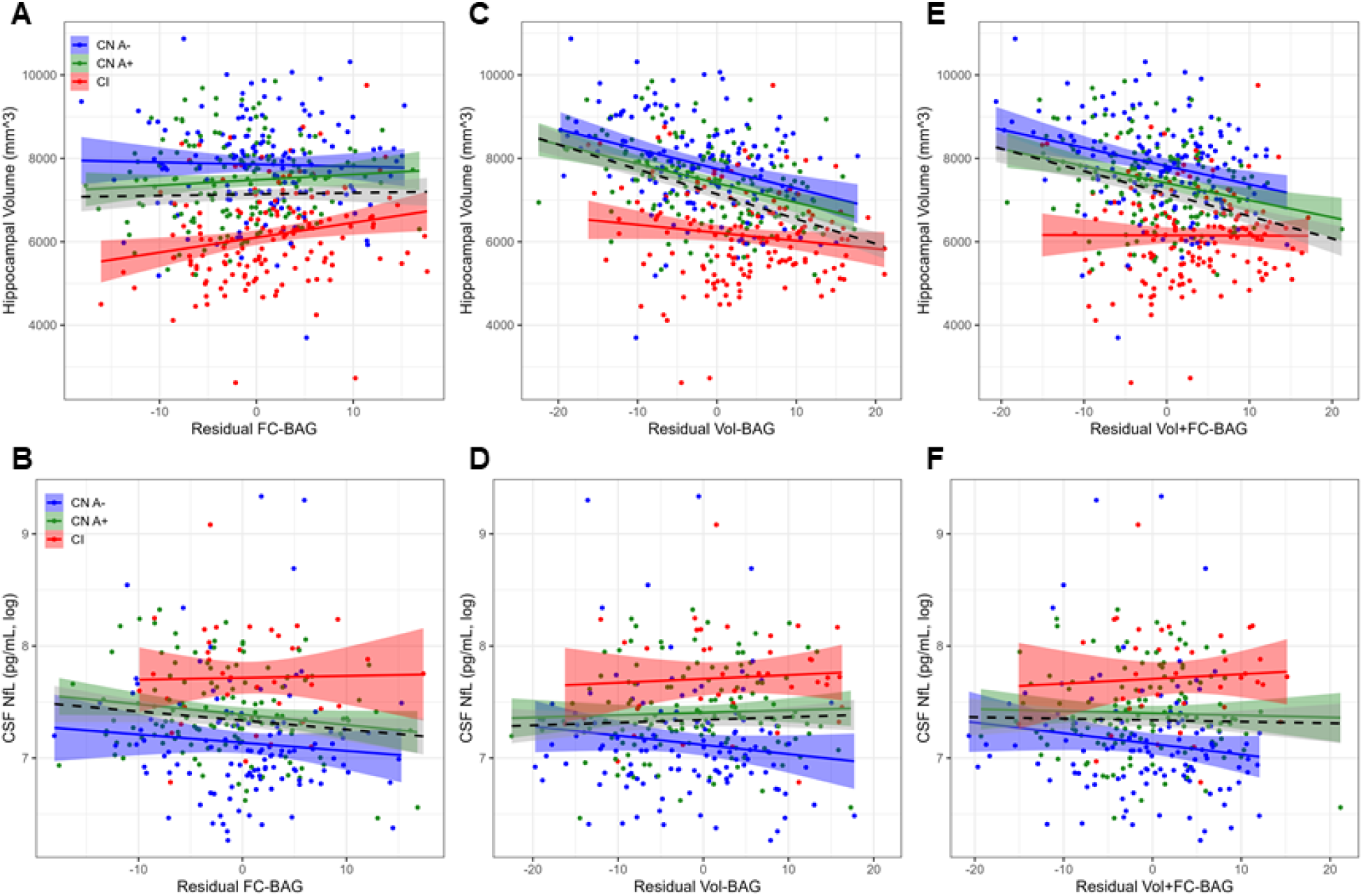
Continuous relationships between neurodegeneration biomarkers and functional connectivity (FC-BAG; A, B), volumetric (Vol-BAG; C, D), and multimodal (Vol+FC-BAG; E, F) brain age gap in the analysis sets. Scatterplots show hippocampal volume (A, C, E) and log-transformed CSF NfL (B, D, F) as a function of residual BAG (controlling for true age) in each group. Colored lines and shaded areas represent group-specific regression lines and 95% confidence regions. Dashed black lines represent main effect regression lines across all groups.

In the volumetric and multimodal models, there were significant main effect relationships, such that greater Vol-BAG (*β* = −53.3,*p* < .001, *η_p_^2^* = 0.193, see Figure 5C) and greater Vol+FC-BAG (*β* = −62.0, *p* < .001, *η_p_^2^* = 0.188, see Figure 5E) were both associated with lower hippocampal volume. There were no significant interactions between the three groups for either model.

In contrast to hippocampal volume, there were no significant main effect relationships between Vol-BAG or Vol+FC-BAG and CSF NfL (see Figure 5D, 5F). However, there were marginal interactions in both estimates, such that the relationships with CSF NfL were more strongly positive in CI participants than in CN A-controls (Vol-BAG: *β* = 0.02, *p* = .057, *η_p_^2^* = 0.013; Vol+FC-BAG: *β* = 0.02, *p* = .083, *η_p_^2^* = 0.011).

### Relationships with Cognition

In the functional connectivity model, there was a significant main effect, such that across all groups, greater FC-BAG was associated with lower cognitive composite score (*β* = −0.02, *p* = .003, *η_p_^2^* = 0.020, see Figure 6A). However, this effect was driven by group differences in both variables, as there were no relationships between FC-BAG and cognition within any of the groups, nor were there any significant interactions.

**Figure 6.**
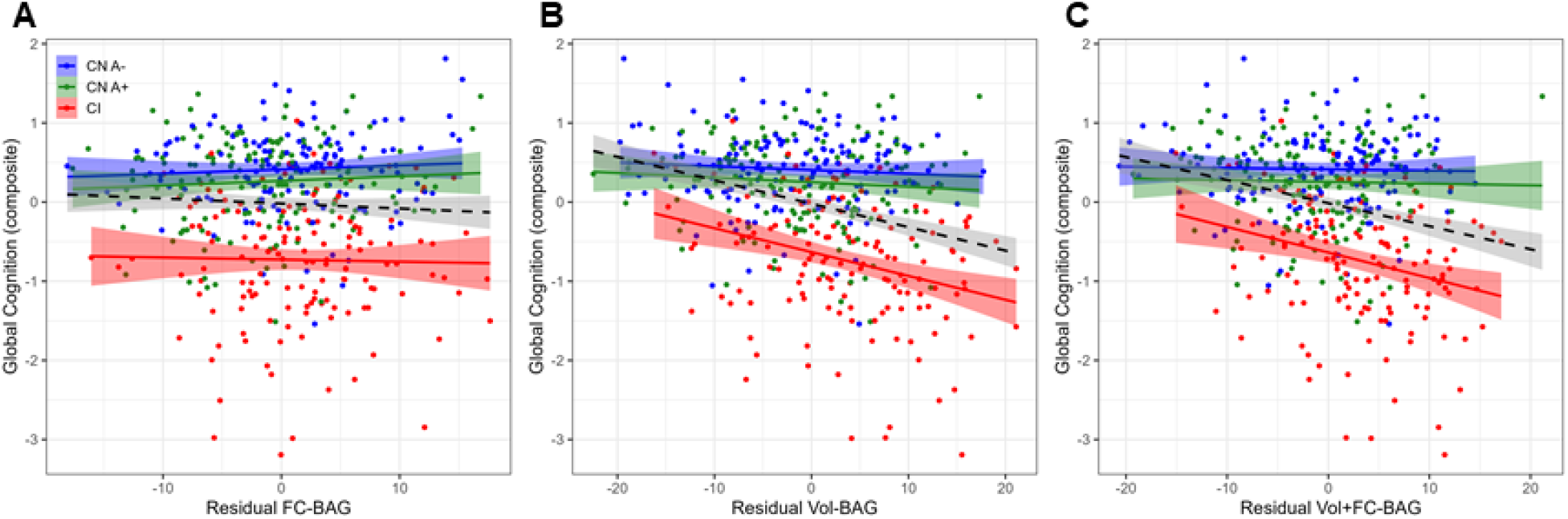
Continuous relationships between global cognition and functional connectivity (FC-BAG; A), volumetric (Vol-BAG; B), and multimodal (Vol+FC-BAG; C) brain age gap in the analysis sets. Scatterplots show global cognition as a function of residual BAG (controlling for true age) in each group. Colored lines and shaded areas represent group-specific regression lines and 95% confidence regions. Dashed black lines represent main effect regression lines across all groups.

In the volumetric model and multimodal models, there were significant main effects, such that greater Vol-BAG (*β* = −0.03, *p* < .001, *η_p_^2^* = 0.104, see Figure 6B) and greater Vol+FC-BAG (*β* = −0.03, *p* < .001, *η_p_^2^* = 0.100, see Figure 6C) were both associated with lower cognitive composite scores. Both effects were further characterized by significant interactions such that the negative associations were observed in the CI participants, but not in the other groups (Vol-BAG: *β* = −0.03, *p* < .001, *η_p_^2^* = 0.045; Vol+FC-BAG: *β* = −0.04, *p* < .001, *η_p_^2^* = 0.048).

## Discussion

We first found that machine learning models successfully predicted age when trained on FC, volumetric MRI, and multimodal datasets. As expected, the volumetric model predicted age with greater accuracy than the FC model, but the multimodal model outperformed both unimodal models. Second, BAG estimates from all models were significantly elevated in CI participants compared to CN controls. BAG estimates in the FC and multimodal models were marginally reduced in preclinical AD participants with elevated amyloid and pTau, but no preclinical group differences were observed in the volumetric model. Third, interactive relationships were observed, such that greater BAG was associated with greater continuous AD biomarker load in CI, but not in CN, participants. Specifically, in the FC model, these interactive effects were only marginally observed in relation to CSF pTau/Aβ40. However, in the volumetric model, these interactions were significantly observed in relation to CSF pTau/Aβ40 and tau PET and marginally with CSF NfL. In the multimodal model, these same interactions were also observed in addition to a marginal interaction with amyloid PET. Finally, regarding cognitive relationships, similar interactive patterns were observed, such that in CI participants, greater BAG estimates from volumetric and multimodal models were associated with lower cognitive performance, however this relationship was not observed in the FC model.

### Predicting Brain Age with Multiple Modalities

We found that a GPR model trained on volumetric MRI features predicted chronological age in a CN adult sample with an *MAE* of about 6 years. This level of performance is comparable to other structural models, which have reported *MAEs* as low as 3 to 5 years (3,13,18,25,72–74). As previously reported (29), the FC-trained model predicted age with an *MAE* of about 8 years, again consistent with previous FC models, which have achieved *MAEs* from 5 to 11 years (25–28). Our observation that volumetric MRI outperformed FC in age prediction is also consistent with previous direct comparisons between modalities (24–26).

Importantly, however, there was only a modest positive correlation between FC and volumetric BAG estimates, after correcting for age-related biases, suggesting that functional and structural MRI capture distinct age-related signals. Indeed, the multimodal model outperformed both unimodal models by integrating these complementary signals. These observations, again, are consistent with other recent reports of multimodal age prediction models (24–27). Future models may improve age prediction accuracy by combining data from volumetric, FC, and/or other neuroimaging modalities, several of which may be available in typical MRI sessions of multiple sequences.

### Brain Age Gap as a Marker of Cognitive Impairment

Volumetric BAG was elevated by 4.66 years in CI participants compared to CN controls. This effect is comparable to previous structural age prediction models, demonstrating elevations in AD and MCI samples between 5 and 10 years (3,4). As previously reported, FC BAG was also elevated in CI participants, but to a relatively smaller extent, i.e. 3.75 years (29). The multimodal BAG was similarly elevated in CI participants by 4.73 years. Thus, each model is clearly sensitive to group differences in AD status at the symptomatic stage.

Consistent with one previous report (17), we demonstrated that within the CI participants, BAG estimates were related to individual differences in AD biomarkers and cognitive function. These effects were most pronounced in the volumetric model, which showed relationships with tau biomarkers and cognition in the CI participants, and the multimodal model, which showed relationships with tau, cognition, and amyloid PET. Thus, age prediction models that include volumetric MRI (including unimodal and multimodal approaches) may be useful in tracking AD pathological progression and cognitive decline within the symptomatic stage of the disease.

### Brain Age Gap as a Marker of Preclinical AD

We found that volumetric BAG did not differ with the presence or absence of preclinical AD biomarkers (amyloid and pTau). In cognitively normal participants, volumetric BAG estimates did not significantly associate with individual differences in any AD biomarkers, except for hippocampal volume. However, it is critical to note that hippocampal volume was indeed an input feature included in the volumetric GPR model, and thus, this finding is not surprising. Overall, although volumetric and multimodal BAG estimates track well with some biomarkers of AD pathophysiology, as previously reported (17), our novel results suggest that these relationships are not observed until the symptomatic phase of the disease, at which point structural changes become more apparent.

As we have previously reported (29), FC BAG was surprisingly lower in preclinical AD participants (particularly A+T+) compared to biomarker-negative controls. Extending beyond this group difference, we now also note that FC-BAG was negatively correlated with amyloid PET in preclinical AD participants. The combined reduction of FC-BAG in the preclinical phase and increase in the symptomatic phase suggest a biphasic functional response to AD progression, which is partially consistent with some prior suggestions (75–79) (see [29] for a more detailed discussion). Notably, this pattern was also marginally observed in the multimodal model, which included FC feature inputs.

Interpretation of this biphasic pattern is still unclear, although the present results provide at least one novel insight. Specifically, one potential interpretation is that the “younger” appearing FC pattern in the preclinical phase may reflect a compensatory response to early AD pathology (80). This interpretation leads to the prediction that reduced FC-BAG should be associated with better cognitive performance in the preclinical stage. However, this interpretation is not supported by the current results, as FC-BAG did not correlate with cognition within any of the analysis samples.

Alternatively, pathological AD-related FC disruptions may be orthogonal to healthy age-related FC differences, as supported by our previous observation that age and AD are predicted by mostly non-overlapping FC networks (29). For instance, the “younger” FC pattern in preclinical AD may be driven by hyper-excitability in the preclinical stage (81,82). Finally, this effect may simply be spuriously driven by sample-specific noise and/or statistical artifacts related to regression dilution and its correction (70). Hence, future studies should attempt to replicate these results in independent samples and further test potential theoretical interpretations.

### Brain Age Gap as a Marker of Cognition

Although FC-BAG was not associated with individual differences in a global cognitive composite within any of our analysis samples, greater volumetric and multimodal BAG estimates were associated with lower cognitive performance within the CI participants. Hence, these estimates may be sensitive markers of cognitive decline in the symptomatic phase. This finding is consistent with previous reports that other structural brain age estimates are associated with cognitive performance in AD (24), Down syndrome (16), HIV (21,32), as well as cognitively normal controls (83).

### Limitations

The training sets included MRI scans from a range of sites, scanners, and acquisition sequence parameters, which may introduce noise and/or confounding variance into MRI features. We attempted to mitigate this problem by: (1) including only data from Siemens 3T scanners with similar protocols; (2) processing all MRI data through common pipelines and quality assessments; and (3) harmonizing across sites and scanners with ComBat (60).

Although we took appropriate steps to detect and control for AD-related pathology in the CN training sets, we were unable to control for other non-AD pathologies, e.g., Lewy body disease, TDP-43, etc., which may be present.

pTau positivity was determined in a data-driven method using the CSF pTau/Aβ40 ratio. Tau PET was only available in a small subset of participants. Future studies might improve upon this approach by using an *a priori* validated threshold of positivity and/or a larger tau PET sample.

Finally, although the Ances lab controls were relatively diverse, participants in other samples were mostly white and highly educated. Hence, these models may not be generalizable to broader samples. Future models would benefit by using more representative training samples.

### Conclusions

We compared three MRI-based machine learning models in their ability to predict age, as well as their sensitivity to early-stage AD, AD biomarkers, and cognition. Although FC and volumetric MRI models were both successful in detecting differences related to healthy aging and cognitive impairment, we note clear evidence that these modalities capture complementary signals. Specifically, FC-BAG was uniquely reduced in preclinical AD participants with elevated amyloid and pTau, although the interpretation of this finding still warrants further investigation. In contrast, volumetric BAG was uniquely associated with biomarkers of AD pathology and cognitive function within the CI participants. Finally, the multimodal age prediction model, which combined FC and volumetric MRI, not only further improved prediction of healthy age differences, but also captured both the biphasic AD response observed in the FC model, as well as the sensitivity to biomarkers and cognition in the volumetric model. Thus, multimodal brain age models may be useful maximizing sensitivity to AD across the spectrum of disease progression.

## Acknowledgements

We thank the participants for their dedication to this project, Haleem Azmy, Anna Boerwinkle, and Dimitre Tomov for technical and processing support. This manuscript has been reviewed by DIAN Study investigators for scientific content and consistency of data interpretation with previous DIAN Study publications. We acknowledge the altruism of the participants and their families and contributions of the DIAN research and support staff at each of the participating sites for their contributions to this study. We thank the personnel of the Administration, Biomarker, Biostatistics, Clinical, Genetics, and Neuroimaging Cores of the Knight ADRC, as well as the Administration, Biomarker, Biostatistics, Clinical, Cognition, Genetics, and Imaging Cores of DIAN.

## Appendix 1

**Dominantly Inherited Alzheimer Network consortium: Full Name and Credentials

Sarah Adams, MS; Ricardo Allegri, PhD; Aki Araki,; Nicolas Barthelemy, PhD; Randall Bateman, MD; Jacob Bechara, BS; Tammie Benzinger, MD, PhD; Sarah Berman, MD, PhD; Courtney Bodge, PhD; Susan Brandon, BS; William (Bill) Brooks, MBBS, MPH; Jared Brosch, MD, PhD; Jill Buck, BSN; Virginia Buckles, PhD; Kathleen Carter, PhD; Lisa Cash, BFA; Charlie Chen, BA; Jasmeer Chhatwal, MD, PhD; Patricio Chrem Mendez, MD; Jasmin Chua, BS; Helena Chui, MD; Laura Courtney, BS; Carlos Cruchaga, PhD; Gregory S Day, MD; Chrismary DeLaCruz, BA; Darcy Denner, PhD; Anna Diffenbacher, MS; Aylin Dincer, BS; Tamara Donahue, MS; Jane Douglas, MPh; Duc Duong, BS; Noelia Egido, BS; Bianca Esposito, BS; Anne Fagan, PhD; Marty Farlow, MD; Becca Feldman, BS, BA; Colleen Fitzpatrick, MS; Shaney Flores, BS; Nick Fox, MD; Erin Franklin, MS; Nelly Joseph-Mathurin, PhD; Hisako Fujii, PhD; Samantha Gardener, PhD; Bernardino Ghetti, MD; Alison Goate, PhD; Sarah Goldberg, MS, LPC, NCC; Jill Goldman, MS, MPhil, CGC; Alyssa Gonzalez, BS; Brian Gordon, PhD; Susanne Gräber-Sultan, PhD; Neill Graff-Radford, MD; Morgan Graham, BA; Julia Gray, MS; Emily Gremminger, BA; Miguel Grilo, MD; Alex Groves,; Christian Haass, PhD; Lisa Häsler, MSc; Jason Hassenstab, PhD; Cortaiga Hellm, BA; Elizabeth Herries, BA; Laura Hoechst-Swisher, MS; Anna Hofmann, MD; Anna Hofmann,; David Holtzman, MD; Russ Hornbeck, MSCS, MPM; Yakushev Igor, MD; Ryoko Ihara, MD; Takeshi Ikeuchi, MD; Snezana Ikonomovic, MD; Kenji Ishii, MD; Clifford Jack, MD; Gina Jerome, MS; Erik Johnson, MD, PHD; Mathias Jucker, PhD; Celeste Karch, PhD; Stephan Käser, PHD; Kensaku Kasuga, MD; Sarah Keefe, BS; William Klunk, MD, PHD; Robert Koeppe, PHD; Deb Koudelis, MHS, RN; Elke Kuder-Buletta, RN; Christoph Laske, PhD; Allan Levey, MD, PHD; Johannes Levin, MD; Yan Li, PHD; Oscar Lopez MD, MD; Jacob Marsh, BA; Ralph Martins, PhD; Neal Scott Mason, PhD; Colin Masters, MD; Kwasi Mawuenyega, PhD; Austin McCullough, PhD Candidate; Eric McDade, DO; Arlene Mejia, MD; Estrella Morenas-Rodriguez, MD, PhD; John Morris, MD; James Mountz, MD; Cath Mummery, PhD;N eelesh Nadkarni, MD, PhD; Akemi Nagamatsu, RN; Katie Neimeyer, MS; Yoshiki Niimi, MD; James Noble, MD; Joanne Norton, MSN, RN, PMHCNS-BC; Brigitte Nuscher,; Ulricke Obermüller,; Antoinette O’Connor, MRCPI; Riddhi Patira, MD; Richard Perrin, MD, PhD; Lingyan Ping, PhD; Oliver Preische, MD; Alan Renton, PhD; John Ringman, MD; Stephen Salloway, MD; Peter Schofield, PhD; Michio Senda, MD, PhD; Nicholas T Seyfried, D.Phil; Kristine Shady, BA, BS; Hiroyuki Shimada, MD, PhD; Wendy Sigurdson, RN; Jennifer Smith, PhD; Lori Smith, PA-C; Beth Snitz, PhD; Hamid Sohrabi, PhD; Sochenda Stephens, BS, CCRP; Kevin Taddei, BS; Sarah Thompson, PA-C; Jonathan Vöglein, MD; Peter Wang, PhD; Qing Wang, PhD; Elise Weamer, MPH; Chengjie Xiong, PhD; Jinbin Xu, PhD; Xiong Xu, BS, MS;

## Supplementary Material

### Participants

**Figure S1.**
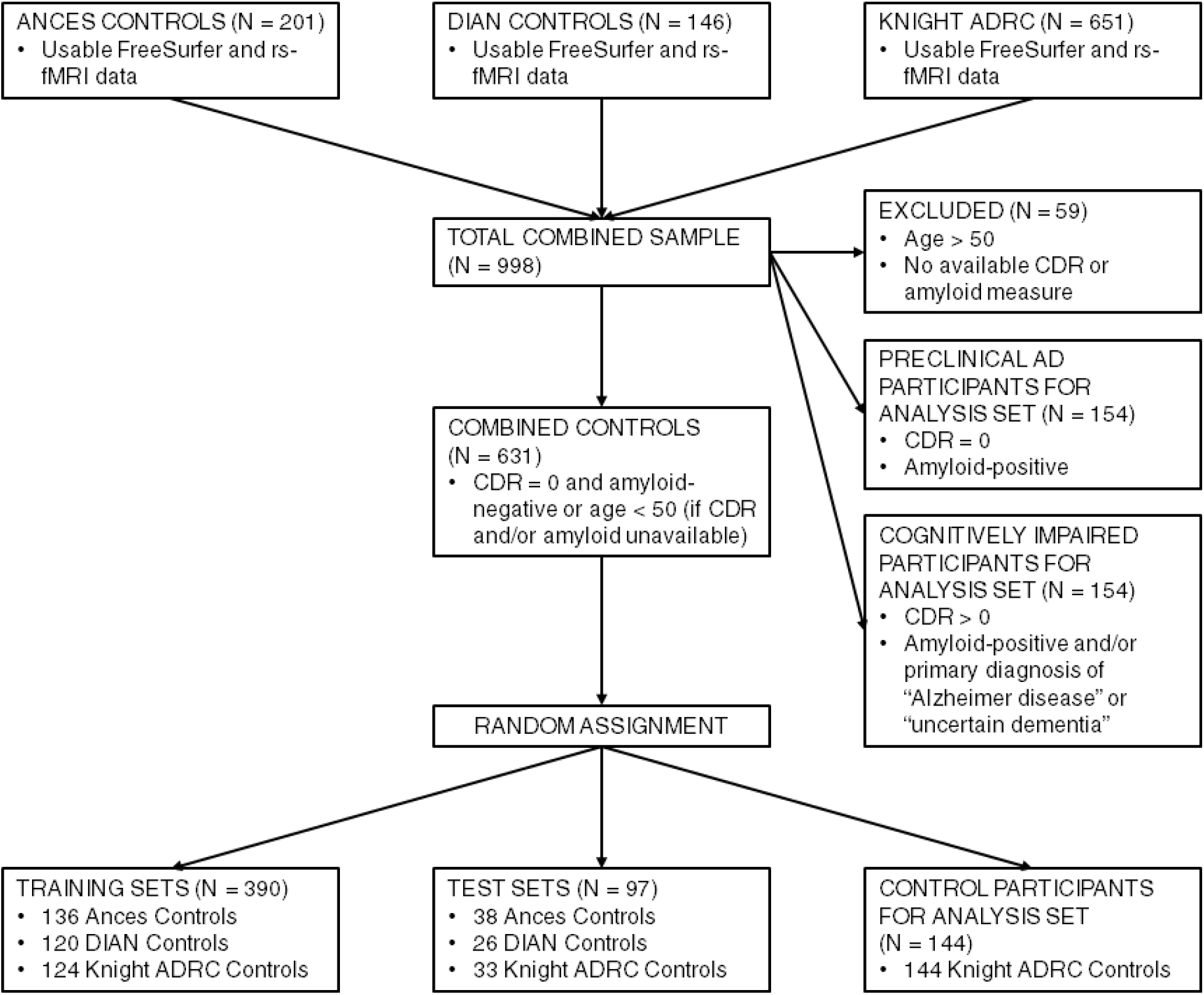
Flow chart of participant inclusion, exclusion, and group assignments.

### GPR Training

**Figure S2.**
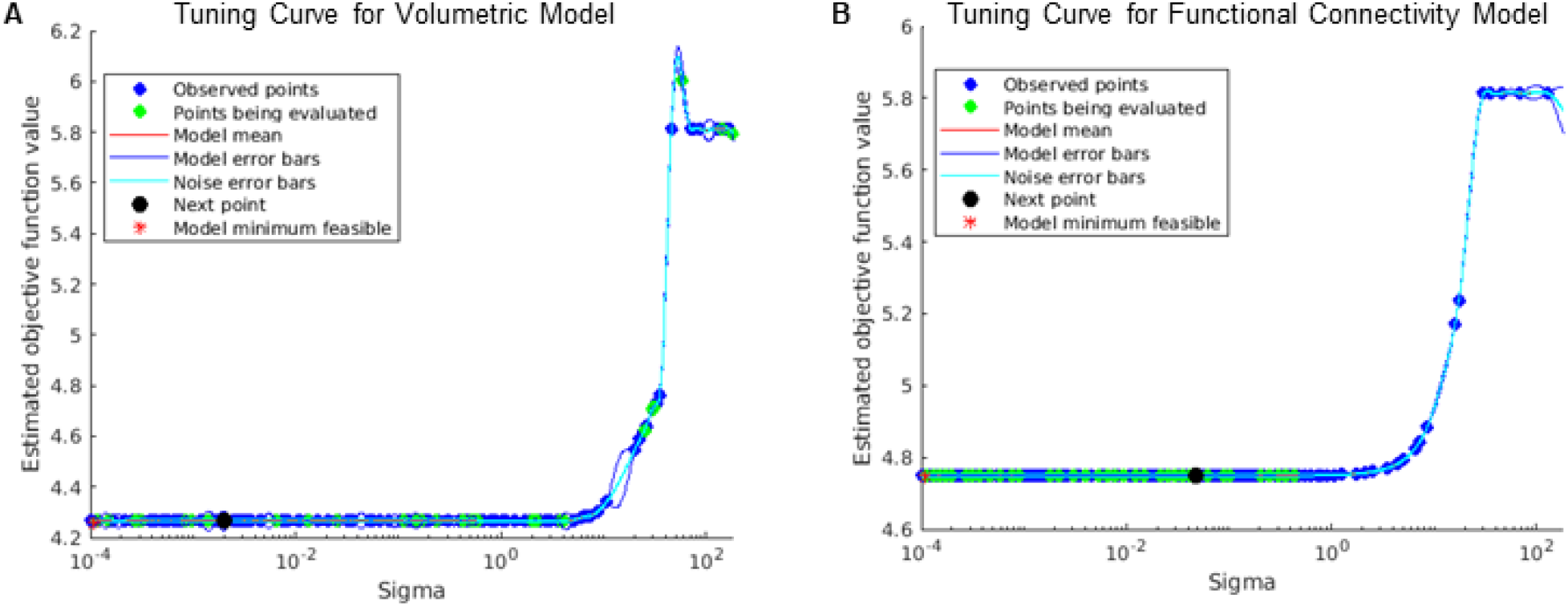
Tuning curves of *σ* hyperparameter in training for volumetric (A) and functional connectivity (B) GPR models.

### Correlation between FC and Vol-Predicted Age

**Figure S3.**
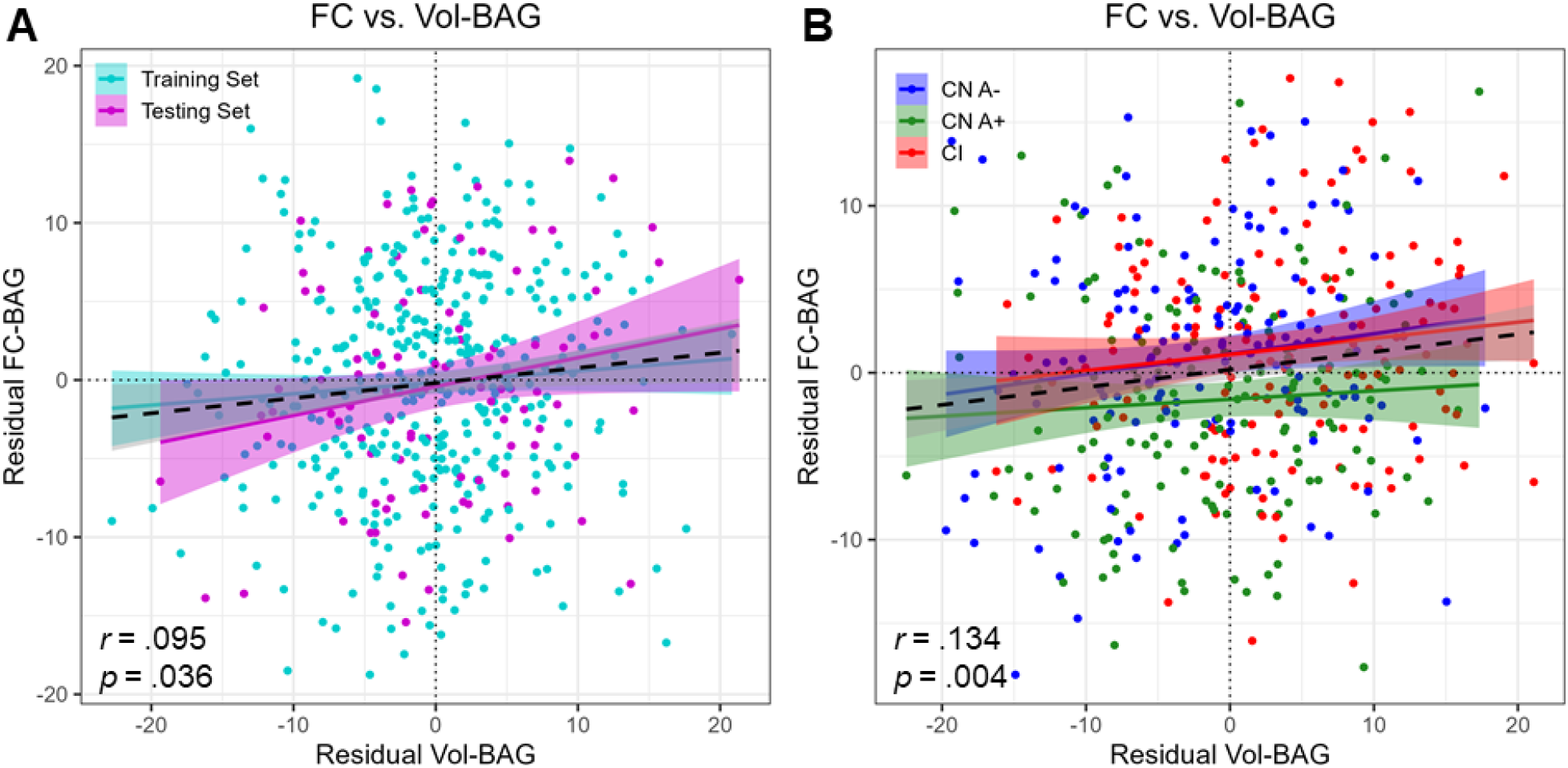
Correlation between Vol-BAG (x axis) and FC-BAG (y axis) estimates in the training and validation sets (A) and analysis sets (B). Both BAG estimates are residualized for age. Dotted black lines represent no difference between predicted and chronological age for each model. Colored lines and shaded areas represent group-specific regression lines and 95% confidence regions. Dashed black lines represent main effect regression lines across all groups.

